# Disc Large Homolog 1 is critical for early microcluster formation and activation in human T cells

**DOI:** 10.1101/2021.05.24.445526

**Authors:** Guha June, Chari Raj

## Abstract

T cell activation by antigen involves multiple sequential steps, including TCR-microcluster (MC) formation, immunological synapse formation and phosphorylation of mediators downstream of the TCR. The adaptor protein Disc Large Homolog 1 (DLG1) is known to regulate proximal TCR signaling and, in turn, T cell activation, acting as a molecular chaperone that organizes specific kinases downstream of antigen recognition. Here, we report using knockdown and knockout studies in human T cells that DLG1 functions even earlier to regulate T cell activation by promoting TCR-MC formation. Moreover, we found that DLG1 can act as a bridge between the TCR-ζ chain and ZAP70 while inhibiting binding of the phosphatase SHP1 to TCR-ζ. Together, these effects drive dysregulation of T cell activation in DLG1-deficient T cells.

## Introduction

The adaptive immune response is critically dependent on T cell activation and plays an important pathophysiologic role in multiple processes, from pathogen clearance to development of autoimmune disease. T cell activation starts with recognition of cognate peptide-MHC (pMHC) complexes on antigen-presenting cells by the TCR on T cells. TCR activation triggers a downstream signal transduction cascade that equips the cell with proper effector function, thus dictating the nature of the immune response [1].

After antigen (Ag) engagement, Lck (a SRC family kinase) is recruited to the TCR-CD3 complex, where it becomes autophosphorylated and activated to then phosphorylate the immunoreceptor tyrosine-based activation motifs (ITAMs) of the CD3 zeta chains. This, in turn, opens up a docking site for ZAP70 (a SYK family kinase) which gets recruited to the ζ chain of the TCR-CD3 complex via its SRC-homology 2 (SH2) domain [2]. Lck activates TCR-bound ZAP70, which in turn phosphorylates its downstream substrate LAT (linker for activation of T cell). Subsequently, LAT recruits several signaling molecules to form a multiprotein complex, termed the LAT signalosome [3]. The LAT signalosome propagates additional signal transduction events, leading to further activation of transcription factors critical for T cell growth and differentiation.

At the cellular level, interaction of the T cell with the APC leads to the formation of the Immunological Synapse (IS) at the interface of the interacting partners. In recent years, it has been shown that Ag-specific activation signals are turned on at TCR-microclusters (TCR-MC) prior to IS formation [4]. TCR-MCs are defined as interacting platforms for the TCR, kinases, phosphatases, and adaptor molecules. TCR-MCs are formed within microseconds of the TCR-pMHC interaction and become building blocks of TCR proximal signaling. They gradually segregate to the cell-cell interface along with adhesion molecules to form the Supra molecular activation complex (SMAC), which consists of a central region known as the c-SMAC and a peripheral adhesion molecule (LFA-1)-rich region known as the p-SMAC. Studies of very early activation events using Ag-specific T cells and a planar bilayer containing specific Ag peptide-MHC has revealed that TCR clustering starts immediately after its interaction with the p-MHC, an event happening much earlier than IS formation. In previous studies with Jurkat cells, TCR stimulation by immobilized anti-CD3 mAb also resulted in induction of signaling cascades [5].

Maintenance of a mature IS requires strong and continuous TCR engagement and downstream signal transduction, and this leads to maximal T cell proliferation and effector function. Adapter or scaffold proteins act as molecular chaperones linking the T cell receptor to proximal signaling moieties, thus controlling the outcome of receptor engagement [6]. The membrane-associated guanylate kinases (MAGUKs) are scaffold proteins known to have critical roles in synaptic development [7]. Members of the MAGUK protein family contain PDZ domains, SH3 (src-homology 3) domains and a guanylate kinase-homologous region (GUK) and often regulate T cell polarity and morphology during migration and immunological synapse formation [8]. Two crucial MAGUKs implicated in T cell function are Carma1 and DLG1 [9, 10]. The human homolog of Drosophila Disc Large, DLG1 translocates to the IS, interacts with Lck and forms a molecular complex with ZAP70 and Wasp [11]. It also coordinates alternative p38 kinase activation and NFAT-driven T cell signaling [12]. Regulatory T cell activation is also influenced by DLG1 [13]. Three distinct Dlg1 deficient mouse strains have been developed, one conditional and two non-conditional. However, no defects in T cell development, morphology, migration, signaling and/or proliferation were observed, suggesting that compensatory mechanisms at different stages of T cell development may mask DLG1 function in T cells in vivo [14]. In the present study, we aimed to further understand the role of DLG1 in proximal human T cell signaling, particularly at the very early stage of antigen recognition and TCR-MC formation.

## Materials and Methods

### Reagents and antibodies

The rabbit antibodies used were as follows: anti-phosphorylated ZAP-70Y319 (2701), anti-phosphorylated PLC-γ1Y783 (2821), anti-phosphorylated LATY171 (3581), anti-phosphorylated ZAP-70Y493 (2704), anti-p44/42 MAPK (anti-ERK1/2; 9101), anti-phosphorylated p44/42 MAPK (anti-phosphorylated ERK1/2; 9102, at Thr202 and Tyr204), anti-ZAP-70 (clone 99F2; 2705), anti-SHP1 (clone-C14H6; 3759), anti-LAT(9166) and anti-p38 MAPK (9212) from Cell Signaling Technologies, California, USA. Anti-CD3-ζ (phosphorylated Tyr83; 68236) from Abcam, Cambridge, UK. The anti-mouse monoclonal antibodies that were used included, anti TCR-ζ (6B10.2;1239), anti-SAP97 (Dlg1; clone 2D11, sc-9961) and anti-Lck (3A5; 433), siRNAs including control siRNA-A (sc-37007), siRNA plasmid against hDlg1 (sc-36452) and hSHP1 (sc-36490)and agarose-conjugated CD3-ζ mouse monoclonal IgG1 (1239-AC) from Santa Cruz, California,USA. Anti-human-CD3 functional grade purified (Clone OKT3,16-0037-85, Clone UCHT1,16-0038-81, Clone HIT3a, 16-0039-81); anti-human-CD28 functional grade purified (clone 28.2, 16-0289-85) from Ebioscience, California, USA. Anti-human CD45 functional grade (Clone HI30,555480), anti-CD247 (P-Y142 CD3, clone K25-407.69, 558402) from BD Pharmingen, California, USA; goat anti-rabbit (11010, Alexa Fluor 564) and goat anti-mouse (21242, Alexa Fluor 647) from Thermo Fisher, Massachusetts, USA; anti β-Actin from Sigma, Missouri, USA.

Anti-P-Y323 p38 from ECM biosciences, Kentucky, USA. Human CD4 recovery column kits (Cedarlane Laboratories Limited, Ontario, Canada). SDS sample buffer was purchased from Quality Biologicals Inc. Maryland, USA, and Restore PLUS western blot Stripping Buffer was from Thermo Scientific Massachusetts, USA.

### Cell line and primary cells

Jurkat cells were obtained from the American Type Culture Collection (ATCC). Jurkat-Zap70 (YFP), a Jurkat line overexpressing YFP-tagged ZAP70 was a kind gift from Dr. Lawrence Samelson (NIH). Human CD4^+^ T cells were isolated from buffy coats of healthy volunteers (NIH Department of Transfusion Medicine) using the human CD4 cell recovery column kit (Cedarlane Laboratories Limited, Ontario, Canada), according to the manufacturer’s instructions.

### Isolation and activation of T cells and immunoblot analysis

Jurkat/ human T cells were incubated with soluble anti-CD3 (OKT3) on ice, followed by cross-linking with anti-mouse-IgG at 37°C for the indicated times. The cells were lysed in 1% Triton-X 100 lysis buffer supplemented with protease and phosphatase inhibitors for 30 min on ice and centrifuged for 10 min at 10,000 r.p.m. The supernatant was boiled in SDS sample buffer for 5 min and subjected to SDS–PAGE, followed by immunoblot analysis.

### Co-Immunoprecipitation

T cells were lysed in ice-cold lysis buffer (150 mM NaCl, 20 mM Tris pH 7.5, 1% Triton X-100 supplemented with 1× protease inhibitor and 1× phosphatase inhibitor; Roche, Basel, Switzerland). Lysates were cleared by centrifugation at 10,000 r.p.m. for 10 min and subjected to immunoprecipitation using agarose-conjugated TCR-ζ or an IgG1 control antibody overnight at 4°C. The beads were washed in ice-cold lysis buffer, and the proteins were eluted by boiling in SDS sample buffer. Finally, the protein was resolved by SDS–PAGE and immunoblotted to analyze the interacting partners.

### Confocal microscopy and image analysis

Cells were allowed to spread on coverslips as described in [5]. In brief, cells were dropped onto polylysine-treated four-chambered glass coverslips coated with stimulatory anti-CD3 (UCHT-1/ HIT3a) or non-stimulatory anti-CD45 (H130) antibodies at 10 μg/ml. Cells were resuspended in warm medium without phenol red, which was supplemented with 25 mM HEPES, pH 7, and dropped at the bottom of the chamber followed by incubation at 37°C for 5 min. The cells were fixed with 2.4% paraformaldehyde for 30 min and permeabilized with Triton X-100. The slides were blocked in blocking buffer (2% goat serum) for 30 min and incubated with primary antibodies for 60 min at the appropriate dilution, followed by secondary antibodies (Alexa-Fluor-568-labeled (1:1,000) or Alexa-Fluor-647-labeled (1:500)) for 45 min. Cells were washed and resuspended in PBS and images were captured using a Leica SP8 laser-scanning confocal microscope using a 63×, 1.4 numerical aperture (NA) objective (Leica Microsystems Inc, Buffalo Grove, IL). 2-to 3-μm z-stacks with a spacing of 0.3 μm were taken of the area contacting the coverslip. Images were processed in Leica AF software and then exported to Adobe Photoshop and Illustrator (Adobe Systems Inc, San Jose CA) to prepare the composite figures.

### siRNA-mediated knockdown

Jurkat or human CD4^+^ T cells were resuspended in 500 μl of antibiotic-free RPMI medium with 10 μl (100 picomoles) each of the validated siRNAs or the control siRNA-A. They were incubated at the room temperature for 5-10 min and they were placed inside a 4-mm cuvette (Bio-Rad). The cuvettes were pulsed using the BTX electroporator (300 V, 10 ms, 960 μF), and the cells were transferred into RPMI medium containing 20% FCS. After 24 hours, the live cells were isolated by Ficoll gradient and resuspended in fresh medium containing 10% FCS. Cells were analyzed for knockdown of DLG1 or SHP1 expression after 48 h by immunoblot analysis.

### CRISPR-Cas9 mediated DLG1 KO and single cell cloning

Guide RNA target sites corresponding to DLG1 were identified using the sgRNA Scorer 2.0 tool (PMID:28146356). Oligonucleotides corresponding to spacer sequences, TCTGGAGAAAAATGCCGGTC and CCAAAAGGTGCAATGCTCTC, were annealed and ligated into the PX458 plasmid backbone using the Enzymatics Rapid T4 ligase. pSpCas9(BB)-2A-GFP (PX458) was a gift from Feng Zhang (Addgene plasmid # 48138http://n2t.net/addgene:48138;RRID:Addgene_48138).

500 ng of cloned plasmid of each guide RNA (1 mg total) were then nucleofected into 250,000 Jurkat cells using the Lonza Nucleofector 4D and the SE nucleofection kit. Electroporation settings used were the manufacturer-suggested settings for the Jurkat-E6. After 72 hours, GFP-expressing cells (transient expression) were sorted and editing efficiency was checked. Single cell sorting was performed on these GFP-positive cells by BD FACSAria Fusion and single cell clones were generated for 2 weeks. Genomic DNA was extracted from single cell clones and NGS sequencing was done to confirm the generation of knock out clones. Putative Dlg1 KO clones were screened for lack of Dlg1 expression by immunoblotting. Further on, these clones were checked for CD3 expression and only high CD3 expressing clones were used for experiments.

Oligonucleotide sequences used for guide RNA cloning are listed below:

Guide1-F CAC CGT CTG GAG AAA AAT GCC GGT C
Guide1-R AAA CGA CCG GCA TTT TTC TCC AGA C
Guide2-F CAC CGC CAA AAG GTG CAA TGC TCT C
Guide2-R AAA CGA GAG CAT TGC ACC TTT TGG C

### Flow Cytometry

CD3 expression was checked using OKT3-APC in BD Calibre. Cells were stained with OKT3-APC in FACS buffer for 30 min at 4°C and analyzed.

### Statistical analysis

Statistical analysis was performed with GraphPad Prism 6 software. Error bars indicate s.e.m. unless otherwise specified. Statistical significance was determined with the paired t-test, and P < 0.05 was considered to be statistically significant.

## Results

### DLG1 knockdown significantly dampens T cell microcluster formation

While the association of DLG1 with the T cell receptor complex has been established, the exact molecular basis and functional significance of this association is still not well understood. In order to interrogate the cellular functionality of DLG1 in T cell synapse formation, we characterized the dynamics of T cell microclusters in Jurkat cells. Using small RNA interference, we demonstrated that DLG1 influences microcluster formation. YFP-tagged ZAP70-expressing Jurkat T cells were dropped onto coverslips coated with stimulatory anti-CD3 or non-stimulatory anti-CD45 specific antibodies. Cells were fixed after 5 min and imaged at the junction of the glass chamber. No distinct ZAP70 clustering could be seen in the cells dropped on the anti-CD45 coated glass chambers. However, when dropped on anti-CD3 coated glass chambers, distinct ZAP70 puncta appeared. Staining with a mAb specific to the TCR zeta chain showed proper colocalization of ZAP70 to TCR-ζ. However, after treatment with siRNA targeting DLG1, puncta intensity was significantly reduced **(Fig 1A and Supplementary Fig 1A, B).** DLG1 is also critical for activation of the T cell alternative p38 pathway [12]. p38 is known to be colocalized with ZAP70 at the immune synapse. [15]. Consistent with ZAP70, phosphorylation of p38 on Y323 was also inhibited under DLG1 knockdown conditions **(Supplementary Fig 1E-H).**

**Figure 1.**
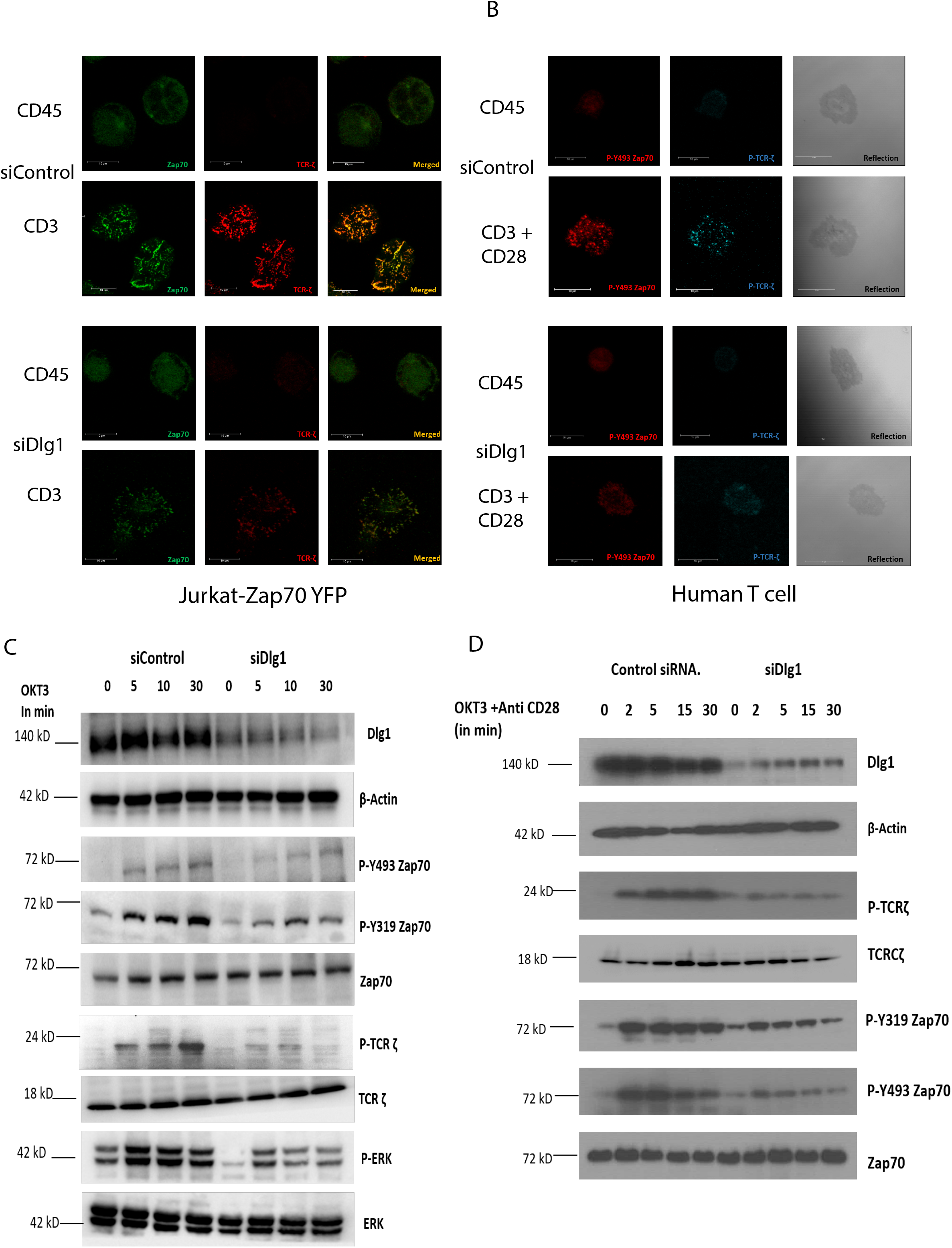
DLG1 knockdown inhibits T cell microcluster formation and subsequent proximal signaling. A and B. Microcluster analysis. Scale bars, 10 μm. Data are representative of three experiments. (A) Jurkat cell analysis. Microclusters of ZAP-70–YFP (green) and phosphorylated TCR-ζ (red) were visualized in immunostained Jurkat cells stably expressing ZAP-70–YFP, pretreated with control or DLG1-specific siRNA. Cells were dropped onto coverslips coated with anti-CD3 (stimulated) or anti-CD45 (unstimulated) and visualized after 5 min. (B) Primary human CD4+ T cells. Microclusters of phosphorylated Y493 ZAP-70 (green) and phosphorylated TCR-ζ (red) were visualized in immunostained human T cells, pretreated with control or DLG1-specific siRNA. Cells were dropped onto coverslips coated with anti-CD3 + anti-CD28 (stimulated) or anti-CD45 (unstimulated) and visualized after 5 min. C and D. Immunoblot analysis of lysates from Jurkat cells (C) and human T cells (D) pretreated with control or DLG1-specific siRNA and then activated with soluble anti-CD3 (monoclonal antibody OKT3) and antibody to mouse immunoglobulin G (IgG), followed by probing for phosphorylated proteins. The membranes were then stripped and reblotted with antibodies to detect total cellular proteins corresponding to the phosphorylated proteins.

Next, we analyzed proximal signaling by evaluating the phosphorylation status of ZAP70 and TCR-ζ. Immunoblot analysis showed that phosphorylation of the activating ZAP70 kinase domain residue Tyr 493 was decreased in the DLG1 KD cells. Similarly, the Tyr 319 residue of ZAP70 and Tyr 83 residue of TCR-ζ also showed reduced phosphorylation. Interestingly, phosphorylation of ERK was diminished by siDLG1 treatment **(Fig 1C).**

Similar results were observed using siRNA treatment in primary human CD4^+^ T cells. Confocal microscopy of freshly isolated human T cells revealed compromised ZAP70 and TCR-ζ activation in DLG1 KD cells **(Fig 1B Supplementary Fig 1C, D).** Concomitantly, diminished phosphorylation of ZAP70 and TCR-ζ was also observed by western blot analysis **(Fig 1D).**

### DLG1 is critical for TCR proximal signaling

In order to reinforce our findings from the DLG1 knockdown experiments, we studied the proximal signaling and microcluster dynamics in DLG1 knockout Jurkat cells using CRISPR-Cas9 gene inactivation. DLG1 KO clones were selected by NGS analysis and confirmed by protein expression studies **(Supplementary Fig S2A,B; Fig S3A).** Clones which showed complete absence of protein were analyzed for TCR expression by flow cytometry **(Supplementary Fig S3B).** The best clone was examined for DLG1 expression and P-Y493 ZAP70 microcluster formation. Confocal images showed compromised microcluster formation in the absence of DLG1 **(Supplementary Fig 3 C-E).**

T cell microcluster dynamics were studied in a similar way in these cells. Wildtype Jurkat cells dropped on stimulatory anti-CD3 coated glass surfaces showed distinct puncta for P-Y493 ZAP70 and P-TCR ζ chain, indicative of microcluster formation at 5 min post activation, as opposed to cells dropped on non-stimulatory anti-CD45 coated glass surfaces. On the contrary, DLG1 KO cells when dropped on stimulatory anti-CD3 coated surfaces showed much reduced intensity of P-Y493 ZAP70 and TCR-ζ clusters **(Fig 2A).**

**Figure 2.**
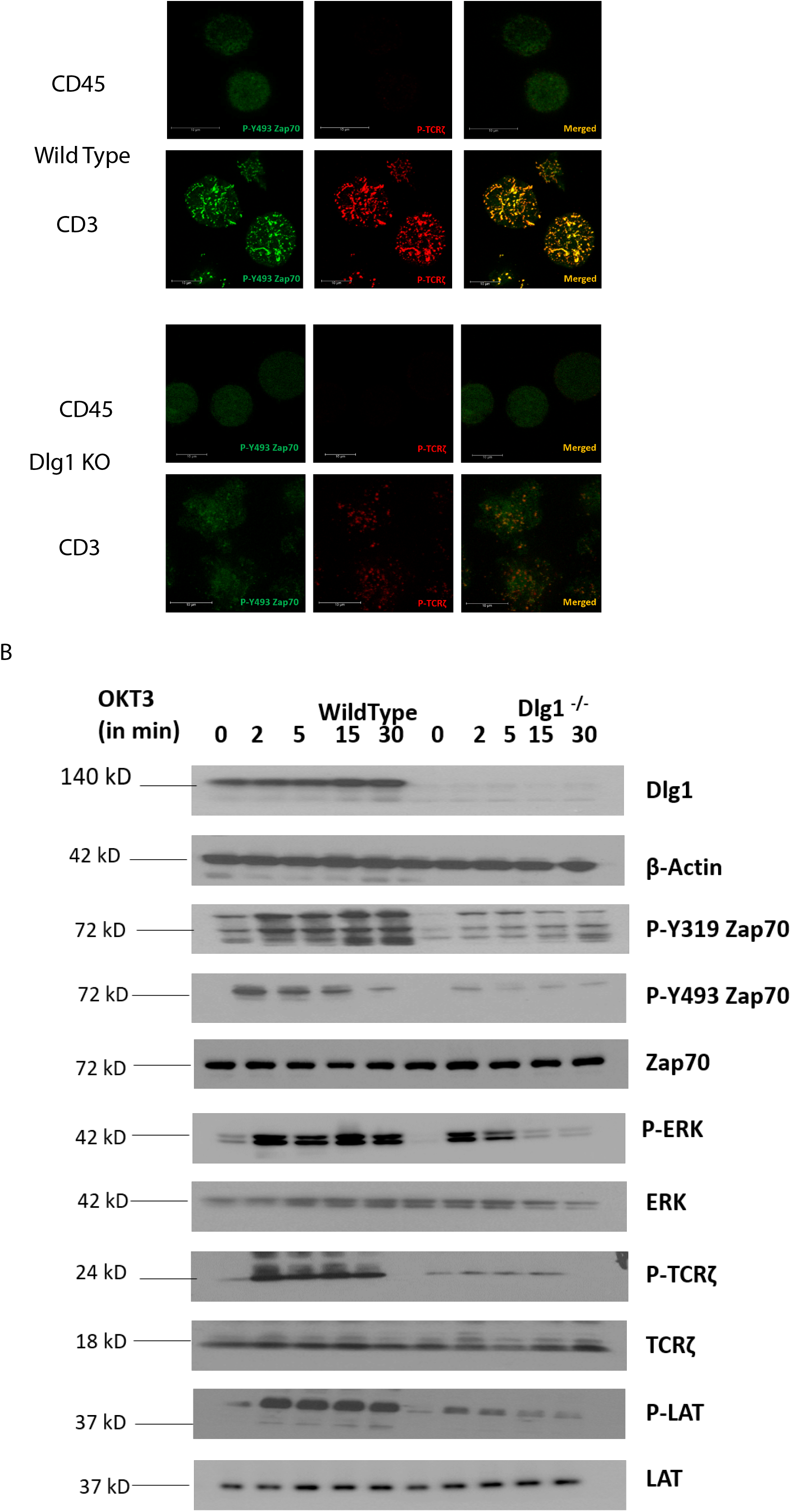
DLG1 knockout inhibits T cell microcluster formation and T cell signaling. A. Microclusters of phosphorylated Y493 ZAP-70(green) and phosphorylated TCR-ζ (red) in immunostained wildtype or DLG1 KO Jurkat cells were visualized 5 min after the cells were dropped onto coverslips coated with anti-CD3 (stimulated) or anti-CD45 (unstimulated) and visualized. Scale bars, 10 μm. Data are representative of three experiments. B. Immunoblot analysis of lysates from wildtype and DLG1 KO Jurkat cells activated with soluble anti-CD3 (monoclonal antibody OKT3) and antibody to mouse immunoglobulin G (IgG). Blots were probed for the indicated phosphorylated proteins, then stripped and reprobed for detection of total cellular proteins corresponding to the phosphorylated proteins.

Downstream signaling was traced by determining the phosphorylation status of the proximal signaling component. Phosphorylation of Y-493 ZAP70 and Y-83 TCR-ζ were significantly reduced as predicted from the quantitative analysis of cluster intensity. Furthermore, phosphorylation of the Y-319 moiety of ZAP70 was also modestly reduced indicative of the compromised activation of ZAP70. Also, activation of LAT and ERK were markedly diminished in DLG1 KO cells **(Fig 2B).**

Overall, the combined evidence from knockdown and knockout experiments supports the notion that DLG1 influences proximal TCR signaling events.

### DLG1 controls the recruitment of ZAP70 to the TCR complex

In order to determine the role of DLG1 in proximal TCR signaling, we performed co-immunoprecipitation (Co-IP) experiments. DLG1 immunoprecipitates were analyzed for TCR proximal proteins. Activation-induced association of TCR-ζ, ZAP70 and LCK with DLG1 was evident in wildtype Jurkat cells **(Fig 3A).** In a reverse Co-IP assay, wildtype and knockout cells were activated with anti-CD3 and cell lysates thereafter were immunoprecipitated with anti-TCR ζ followed by immunoblot analysis. Association of ZAP70, TCR-ζ chain and LCK was observed as early as 5 min after stimulation in wildtype cells. These associated proteins were activated, as determined from analysis of their phosphorylation status. ZAP70 association with the TCR-ζ chain was drastically compromised in DLG1’s absence. Notably, phosphorylation of the TCR-ζ chain was also reduced **(Fig 3B).**

**Figure 3.**
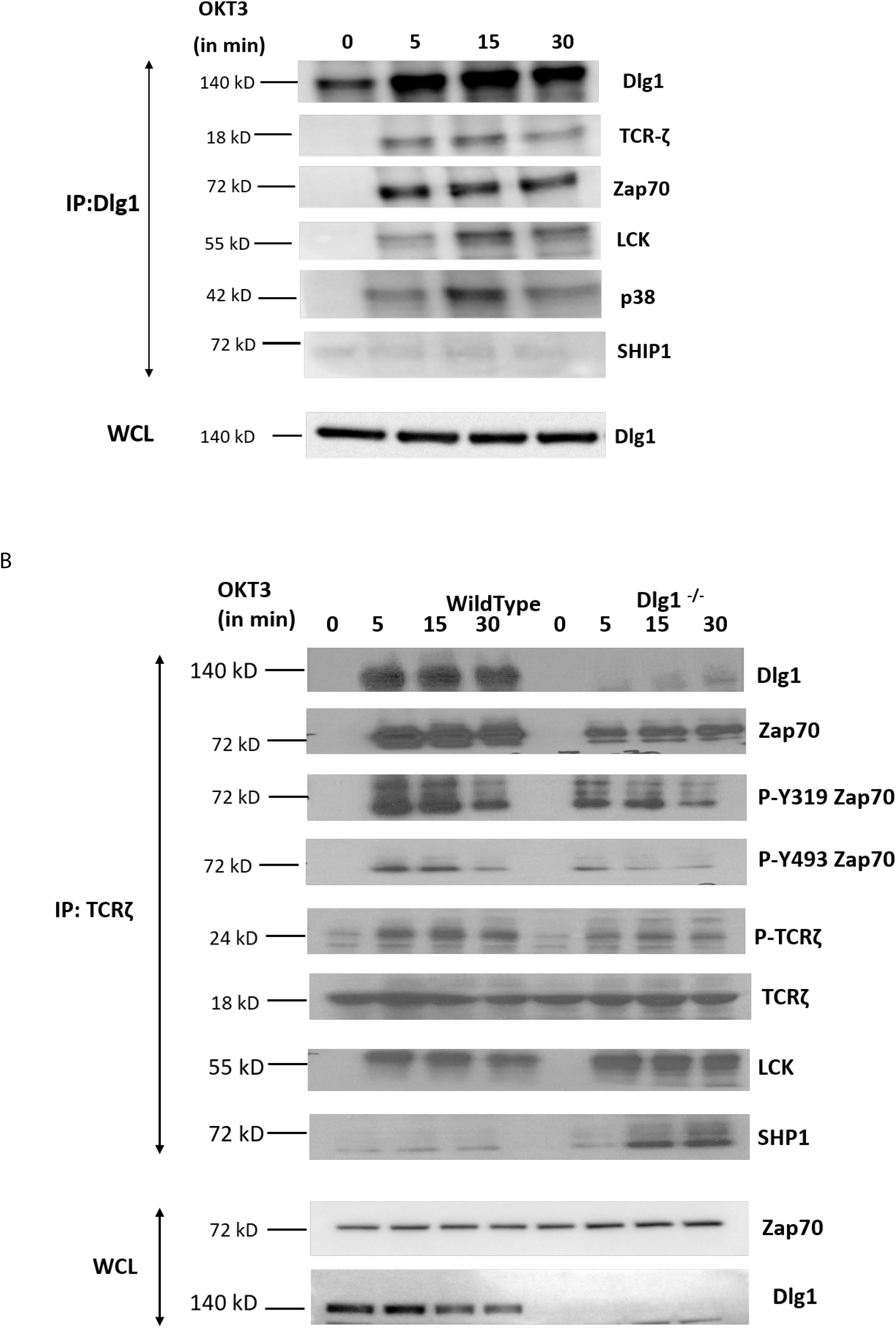
DLG1 is critical for ZAP70 recruitment to the TCR complex. A. Immunoblot analysis (IB) of the indicated co-immunoprecipitated partners after immunoprecipitation (IP) of DLG1 from activated Jurkat cells. Data representative of 2 experiments B. Immunoblot analysis (IB) of the indicated co-immunoprecipitated partners after immunoprecipitation (IP) of TCR-ζ from activated wildtype and DLG1 KO Jurkat cells. Data representative of 2 experiments

ZAP70 is recruited to the TCR complex after the ζ chain is phosphorylated, which opens up a docking site for the protein. As phosphorylation of the ζ chain is compromised in DLG1 KO cells, we examined the status of LCK, the immediate upstream kinase. Surprisingly, LCK recruitment to the complex was unaffected in knockout cells as compared to wildtype cells **(Fig 3B).**

Phosphorylation status of any protein depends upon the balance between phosphorylation and dephosphorylation. Since LCK recruitment was unaffected by DLG1 KO, we examined the status of SHP1, the critical phosphatase for ζ chain dephosphorylation. Interestingly, SHP1 association with the ζ chain was significantly increased in DLG1-deficient cells **(Fig 3B).** Thus, the increased association of SHP1 to the TCR complex under conditions of DLG1 deficiency may be a determinant of dysregulated proximal TCR signaling.

### Enhanced association of SHP1 to the TCR complex compromises proximal TCR signaling

Next, we aimed to evaluate the role of SHP1 in association with DLG1 in T cell signaling. For this, we re-analyzed the microcluster dynamics in Jurkat cells. Wildtype and DLG1 knockout Jurkat cells were treated with an siRNA against SHP1. After 48 hrs, the cells were placed onto an anti-CD3 or anti-CD45 coated surface and microclusters were analyzed. After SHP1 siRNA treatment, wildtype cells showed a modest increase in P-ZAP70 and P-TCR ζ signal intensity. On the other hand, the decreased signal intensity in activated DLG1 KO cells was rescued after SHP1 siRNA treatment **(Fig 4A).**

**Figure 4.**
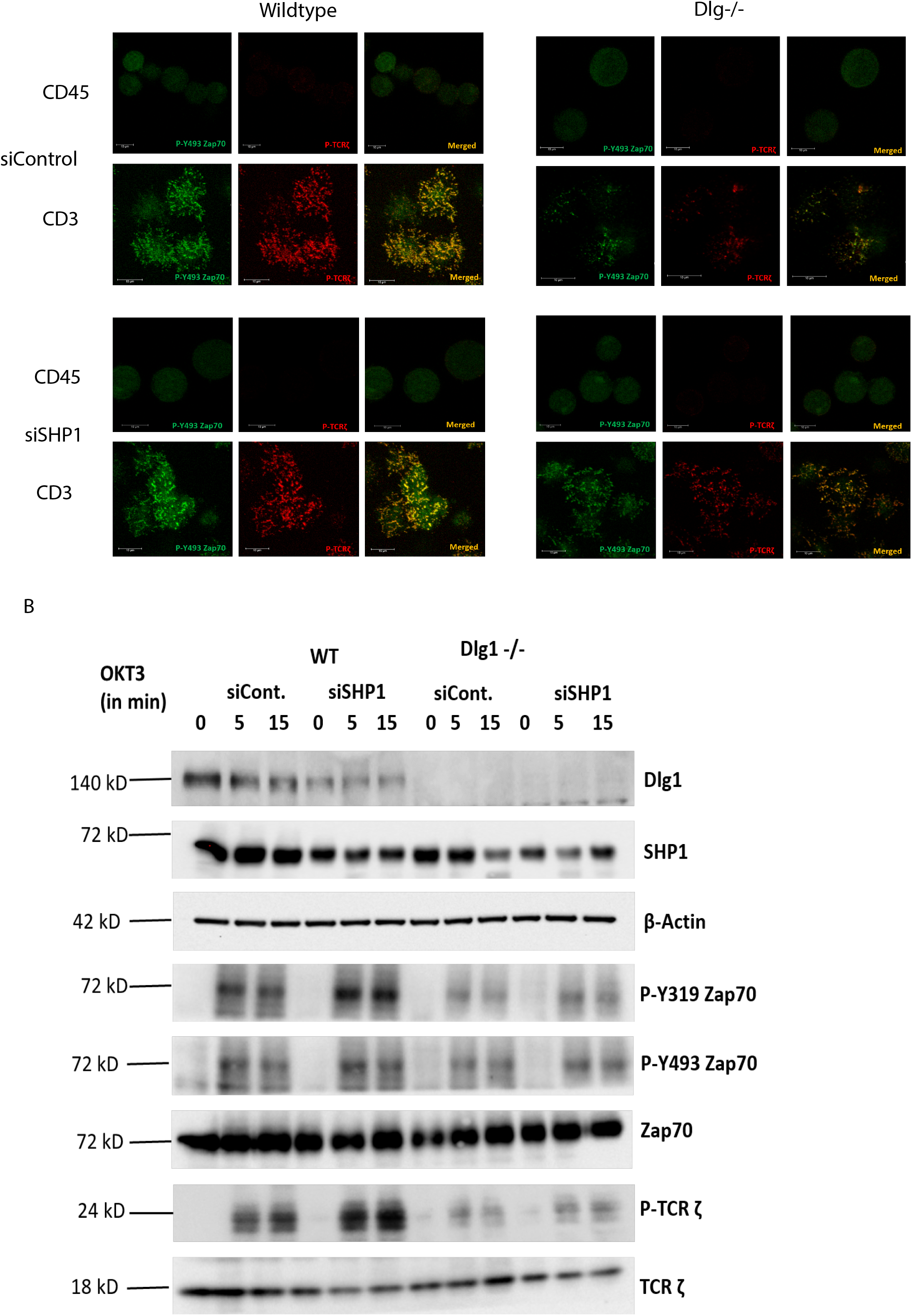
SHP1 knockdown partially rescues the inhibition of T cell microcluster formation and proximal signaling conferred by DLG1 deficiency. A. Microclusters of phosphorylated Y493 ZAP-70 (green) and phosphorylated TCR-ζ (red) were visualized in immunostained wildtype or DLG1 KO Jurkat cells that were pretreated with control or SHP1 specific siRNA. Cells were dropped onto coverslips coated with anti-CD3 (stimulated) or anti-CD45 (unstimulated) and visualized after 5 min. Scale bars, 10 μm. Data are representative of three experiments. B. Immunoblot analysis of lysates from wildtype and DLG1 KO Jurkat cells. Cells were pretreated with control or SHP1-specific siRNA, then activated with soluble anti-CD3 (monoclonal antibody OKT3) and antibody to mouse immunoglobulin G (IgG), followed by probing for phosphorylated proteins. Membrane were then stripped and re-probed for detection of total cellular proteins corresponding to the phosphorylated proteins.

After TCR activation, DLG1 immunoprecipitates contained LCK, Zap70, p38 and TCR-ζ **(Fig 3A).** However, significant association with SHP1 was not observed. The downstream effect of this reversal was determined by phospho-protein analysis. Significant increases in P-Y493 ZAP70 and P-Y319 ZAP70 were observed in DLG1 KO cells after siRNA SHP1 treatment as opposed to control siRNA treatment. Increased P-TCR ζ strongly suggested the loss of phosphatase activity in both wildtype and DLG1 KO cells **(Fig 4B).** siRNA treatment against SHP2 did not show any significant effects **(Supplementary Fig S4A).**

## Discussion

DLG1, a homolog of Drosophila Disc large tumor suppressor protein is a member of the MAGUK (Membrane-associated Guanylate kinase) protein family. It is a scaffolding molecule known to have an important role in the immune synapse, where it organizes signaling components to form synaptic complexes [16]. In epithelial cells, its role is to regulate actin cytoskeleton dynamics and thus to maintain cell polarity [17] [18]. Apart from its role in neuronal and epithelial cells, DLG1 is also known to have a pivotal function in T cell activation [11].

Several adapter proteins have already been implicated in TCR signal transduction. DLG1 is known to translocate to the TCR complex upon receptor engagement. It orchestrates the inter-molecular interaction between signaling components at the TCR such as LCK-ZAP70-WASp [11]. Alternative activation of p38 is mediated by LCK-ZAP70 and an association with DLG1 is evident [12]. TCR-MC which are platforms for signaling cascades includes activated LCK and ZAP-70. ZAP-70 colocalizes with p38 and activates it alternatively by phosphorylating its Tyr323 moiety [15]. Our data demonstrate that the ZAP-70-p38 association is heavily compromised at TCR-MC when DLG1 is knocked down in Jurkat cells and primary human T cells. This proves that DLG1 is working at the immediate phase of T cell activation, even before the formation of the immunological synapse. Consequently, dampening of downstream signaling is observed by Western Blot analysis under conditions of DLG1 deficiency. The decreased phosphorylation of the TCR-ζ chain explains the subsequent reduction of ZAP-70 phosphorylation at the activating residues Tyr493 and Tyr319. Finally, compromised LAT activation leads to defective LAT signalosome formation which acts as a key event in T cell activation. The initial finding with DLG1 knockdown was confirmed by DLG1 knockout experiments. Visualization of the TCR microcluster revealed loss of ZAP-70 and p-TCR-ζ in the microclusters.

Molecular interaction between upstream kinases to their substrates moves as a chain reaction as more and more downstream molecules are involved. Interaction between the ζ chain of CD3 to ZAP70 through its SH2 domain is the key event in T cell signal initiation. In the absence of DLG1, we found reduced interaction of ZAP70 to its immediate partner. This may be explained by nonavailability of the docking site for ZAP70 at the ζ chain. ζ chain is heavily phosphorylated by Lck. However, in the absence of DLG1, a decrease in ζ chain phosphorylation was observed. Interestingly, Lck recruitment to the TCR is intact even in DLG1 deficiency thus ruling out loss of function of the activating kinase. The alternative mechanism which could be proposed is gain of function of deactivating phosphatases. Surprisingly, the SHP1 association with TCR ζ is significantly enhanced in the absence of DLG1, identifying a role in dampening T cell signal initiation.

Studies in T cell proximal signaling show that the opposing actions of protein tyrosine kinases (PTKs) and protein tyrosine phosphatases (PTPs) are always in constant operation and the balance between the two determines the activation/suppression of T cell signaling. Classical PTPs include SHP1 and SHP2, which are SH2 domain-containing phosphatases implicated in T cell signaling [19]. SHP1 is expressed in all hematopoietic cells; however, its expression is low in epithelial cells [20]. In contrast, SHP2 is ubiquitously expressed [19]. In our study, we found that wild type Jurkat cells show enhanced phosphorylation of ZAP70, ERK and TCR ζ when SHP1 is knocked down, identifying a direct effect of SHP1 in suppressing proximal signaling. In Jurkat cells, SHP1 but not SHP2 knockdown partially rescued dampened signaling conferred by DLG1 deficiency. This is in agreement with SHP2 having positive effects or no effect on T cell signaling, although it is found in complex with ITAMs [19].

ZAP70 is known to directly bind to SHP1 [21]; however, the precise molecular mechanism of SHP1 recruitment to the TCR/CD3 complex remains unclear. SHP1 is also known to crosstalk with CEACAM [22] and Themis [23] in order to control the outcome of T cell signaling. Another consideration is the increased phosphorylation status of SHP1. Phosphorylation of Y536 increases SHP1 activity about 4-8-fold, while that of Y564 phosphorylation increases it by 1.6-fold. Whether SHP1 is interacting with other partners more strongly or is selectively phosphorylated to increase its PTP activity is an important question for future research.

Our finding that DLG1 regulates proximal T cell signaling is consistent with the known function of other adaptor proteins in T cell activation. However, the microscopic visualization of the proximal molecules in their activation states provides detailed evidence of molecular interaction at the IS. In a recent publication, the authors showed that SHP1 promotes T cell adhesion by activating the adaptor protein CrkII by dephosphorylating it in the IS [24]. In our study we see a reverse picture, where an adaptor molecule, DLG1, prevents the recruitment of a phosphatase into the IS thereby promoting sustained signaling. In the absence of DLG1 this checkpoint is lost and thus the T cell fails to form an effective MC. Another docking protein, Gab2, is known to be phosphorylated by ZAP70 and to negatively regulate T cell receptor signaling by recruitment of inhibitory molecules [25]. DLG1 works in an opposing manner to inhibit SHP1 interaction with the TCR. In conclusion, DLG1 is a positive regulator of T cell activation, and thus it may be important for immune activation in health and disease.

## Acknowledgement

Jurkat-YFP cells are a kind gift from Dr. Lawrence E Salemson. All the reagents and use of equipment was kindly provided by Dr. Jonathan D Ashwell. Thanks to Dr. Philip M Murphy for carefully editing the manuscript.

## Footnote

The authors declare no conflict of interest.

This work was supported by funding from the Intramural Research Program of the Center of Cancer Research, NCI, NIH and NIAID, NIH.

