## Supplemental File for "Disc Large Homolog 1 is critical for early microcluster formation and activation in human T cells"

Figure S1

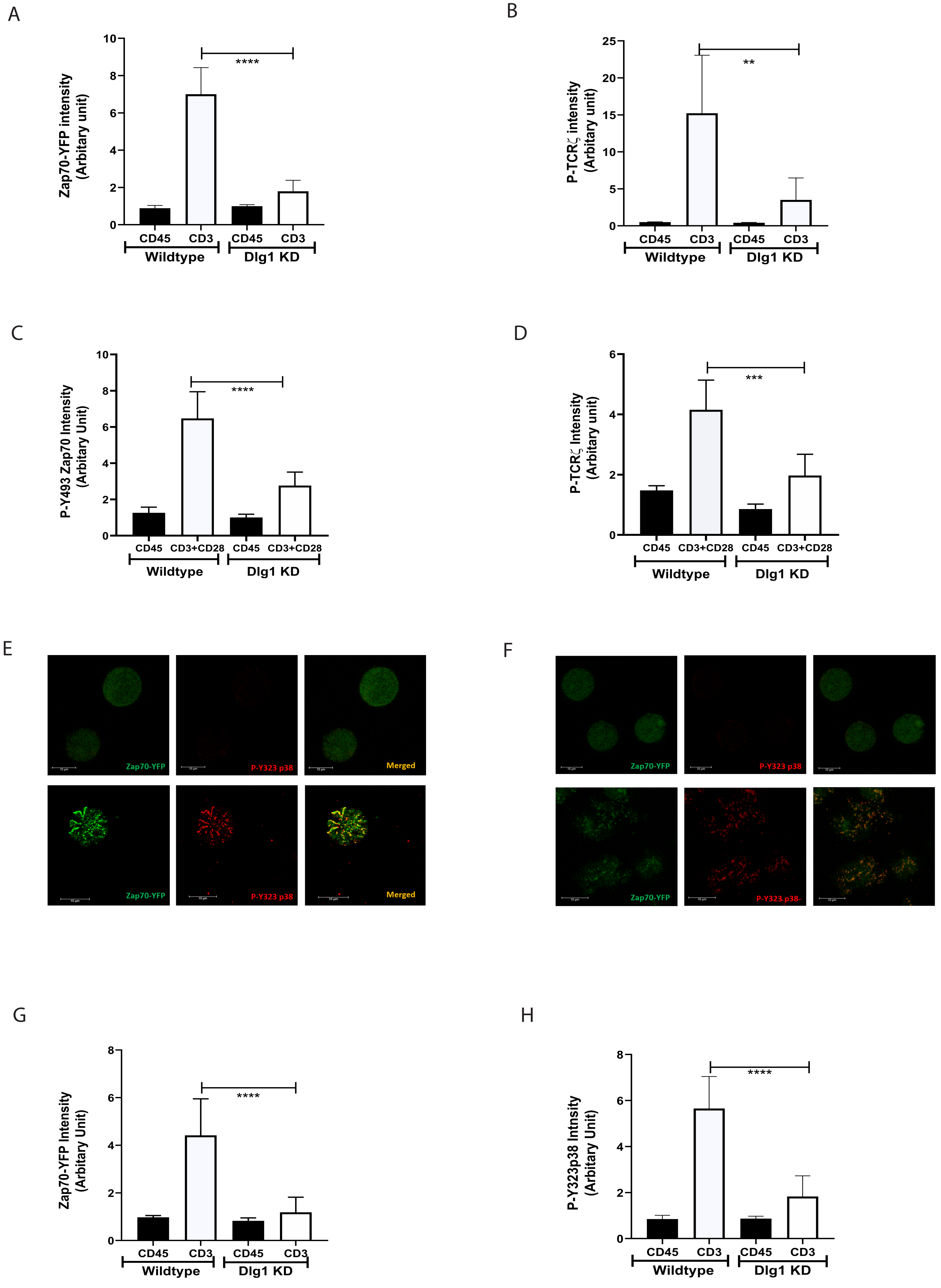

A
Zap70-YFP intensity (Arbitary unit)
CD45
CD3
Wildtype
Dlg1 KD
****
B
P-TCRζ intensity (Arbitary unit)
**
C
P-Y493 Zap70 Intensity (Arbitary Unit)
CD3+CD28
D
P-TCRζ Intensity (Arbitary unit)
***
E
Zap70-YFP
P-Y323 p38
Merged
F
G
Zap70-YFP Intensity (Arbitary Unit)
H
P-Y323p38 Intnsity (Arbitary Unit)

Figure S2

A

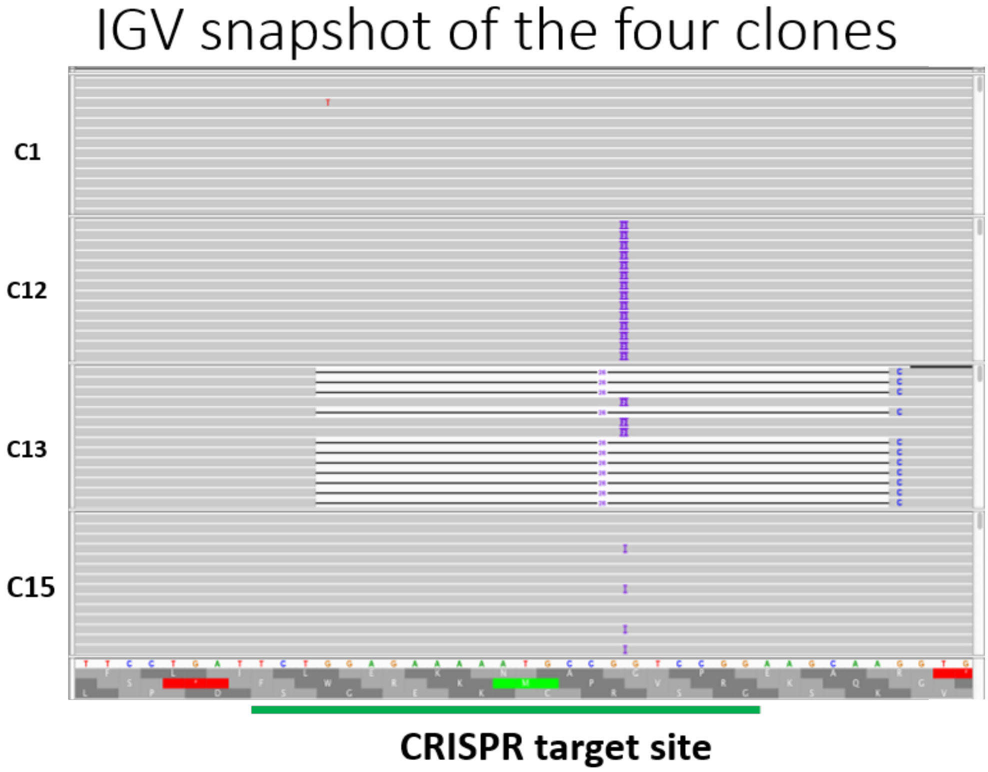


B

Clone

Coding Exon 1

TCTTCCTGAT<u>TCTGGAGAAAA**ATG**CCGGTCCGG</u>AAGCAAGGTGAGAGTTTATTCCGTAACT

C1 Allele 1 TCTTCCTGATTCTGGAGAAAAATGCCGGTCCGGAAGCAAGGTGAGAGTTTATTCCGTAACT

Allele 2 TCTTCCTGATTCTGGAGAAAAATGCCGGTCCGGAAGCAAGGTGAGAGTTTATTCCGTAACT

Allele 1 TCTTCCTGATTCT-------------------------------------------------------------CGTGAGAGTTTATTCCGTAACT

Allele 2 TCTTCCTGATTCTGGAGAAAAATGCCG**TC**GTCCGGAAGCAAGGTGAGAGTTTATTCCGTAACT

Allele 1 TCTTCCTGATTCTGGAGAAAAATGCCG**CTC**GTCCGGAAGCAAGGTGAGAGTTTATTCCGTAACT

Allele 2 TCTTCCTGATTCTGGAGAAAAATGCCG**CTC**GTCCGGAAGCAAGGTGAGAGTTTATTCCGTAACT

Allele 1 TCTTCCTGATTCTGGAGAAAAATGCCG**T**GTCCGGAAGCAAGGTGAGAGTTTATTCCGTAACT

Allele 2 TCTTCCTGATTCTGGAGAAAAATGCCGGTCCGGAAGCAAGGTGAGAGTTTATTCCGTAACT

sgRNA target site underlined

Figure S3

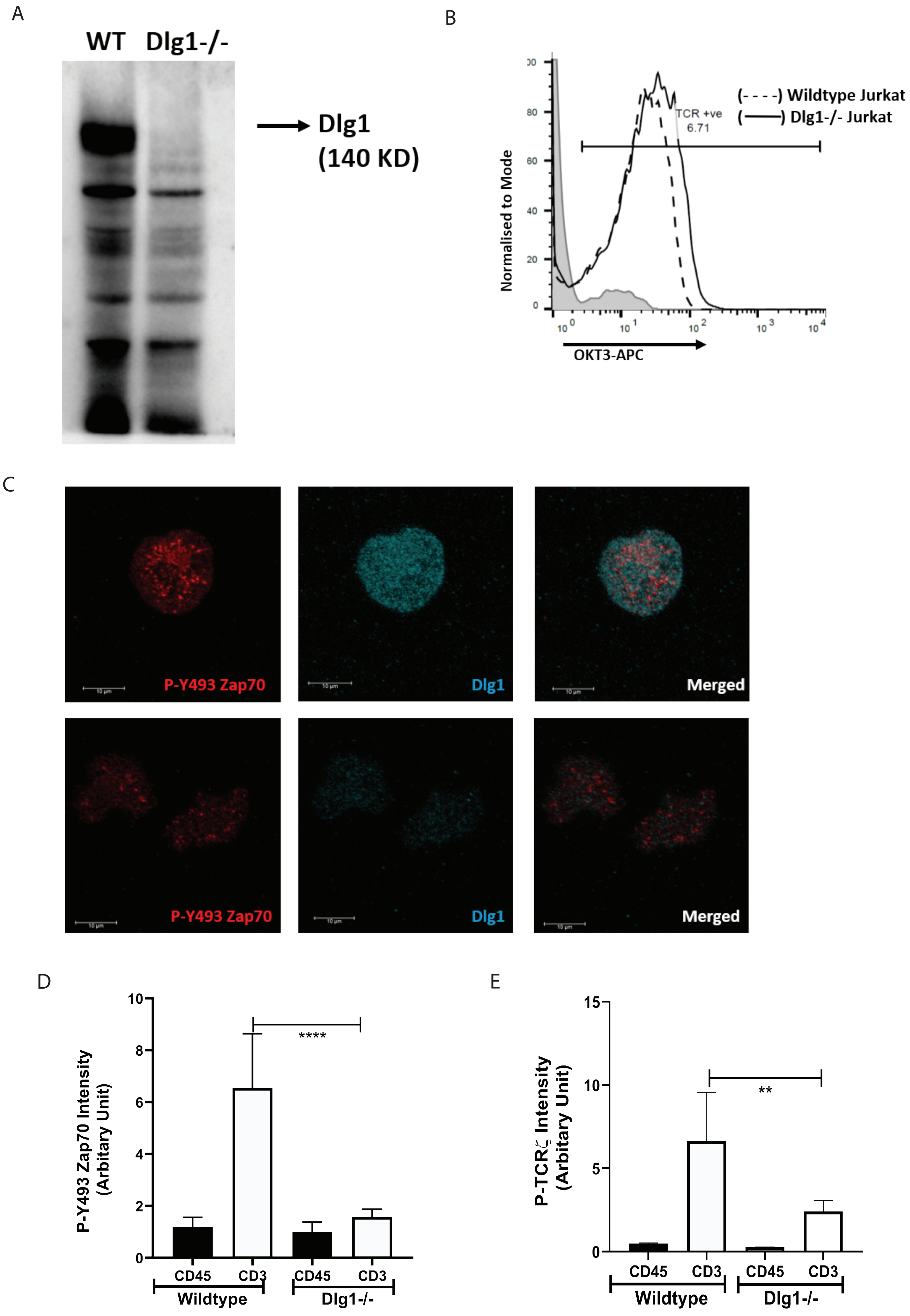
A
WT Dlg1-/-
Dlg1 (140 KD)
B
Normalised to Mode
(- - - -) Wildtype Jurkat
(——) Dlg1-/- Jurkat
TCR +ve 6.71
OKT3-APC
C
P-Y493 Zap70
Dlg1
Merged
P-Y493 Zap70
Dlg1
Merged
D
P-Y493 Zap70 Intensity (Arbitary Unit)
****
CD45 CD3 CD45 CD3
Wildtype Dlg1-/-
E
P-TCRζ Intensity (Arbitary Unit)
**
CD45 CD3 CD45 CD3
Wildtype Dlg1-/-

Figure S4

A

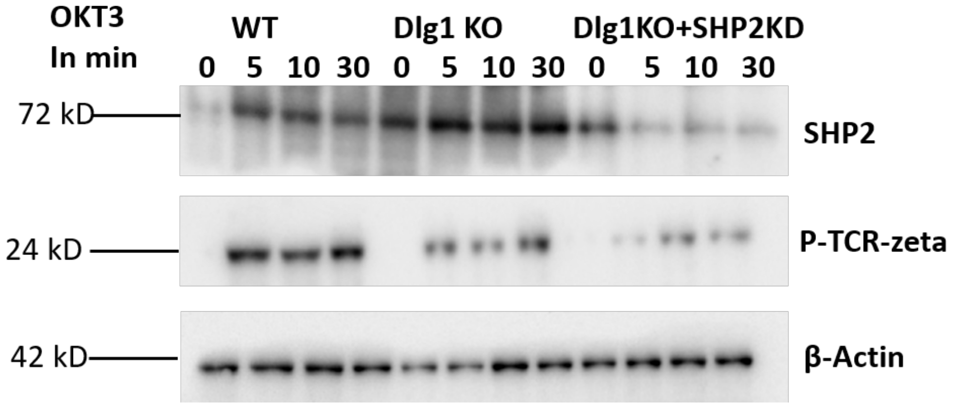


B

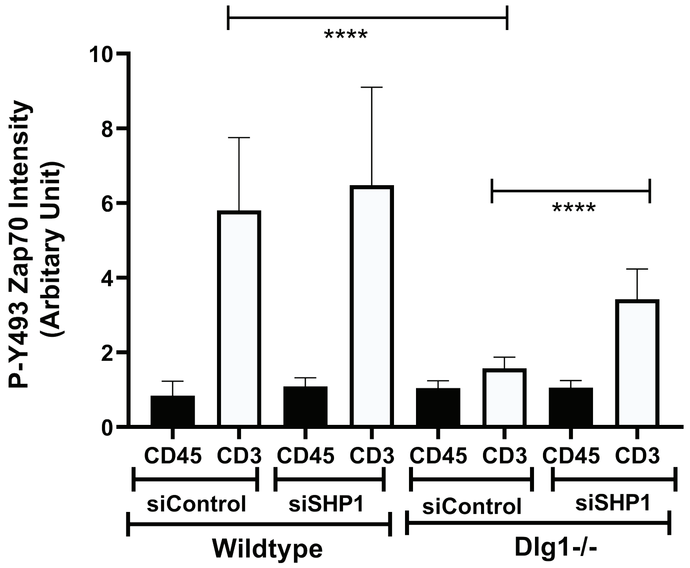


C

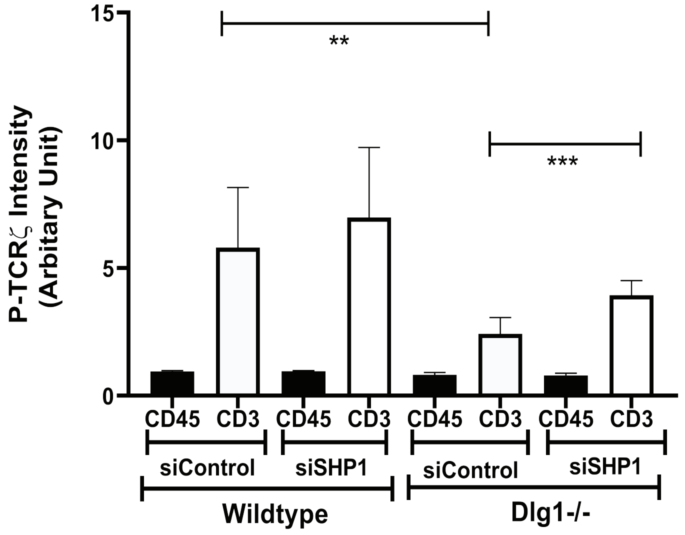

**Supplementary Figure 1** Quantitative analysis of T cell microcluster intensity

A and B. Zap70-YFP and P-TCR ζ intensities were quantified using Image J in immunostained Jurkat cells stably expressing ZAP-70–YFP. Cells were pretreated with control or DLG1-specific siRNA, then dropped onto coverslips coated with anti-CD3 (stimulated) or anti-CD45 (unstimulated). Data are representative of 8 fields from 3 independent experiments.

C and D. Phosphorylated Y493 ZAP-70 and phosphorylated TCR-ζ intensities were quantified using Image J in immunostained human T cells. Cells were pretreated with control or DLG1-specific siRNA, then dropped onto coverslips coated with anti-CD3 + anti-CD28 (stimulated) or anti-CD45 (unstimulated). Data are representative of 8 fields from 3 independent experiments.

E and F. Microclusters of ZAP-70–YFP (green) and phosphorylated Y323 p38 (red) were imaged in immunostained Jurkat cells stably expressing ZAP-70–YFP. Cells were pretreated with control or DLG1 specific siRNA, then dropped onto coverslips coated with anti-CD3 (stimulated) or anti-CD45 (unstimulated) and visualized after 5 min. Scale bars, 10 µm. Data are representative of three experiments

G and H. ZAP70-YFP and P-Y323 p38 intensities were quantified using Image J in immunostained Jurkat cells stably expressing ZAP-70–YFP. Cells were pretreated with control or DLG1-specific siRNA, then dropped onto coverslips coated with anti-

CD3 (stimulated) or anti-CD45 (unstimulated). Data are representative of 8 fields from 3 independent experiments.

**Supplementary Figure 2** Sequence analysis of DLG1 KO clones

A. IGV snapshot of DLG1 KO clones

B. Allele analysis of Clonal populations

**Supplementary Figure 3** Characteristic analysis of DLG1 KO cells

A. Immunoblot of DLG1 from wildtype and DLG1 KO Jurkat cell

B. TCR-ζ expression in wildtype and DLG1 KO Jurkat cell

C. Microcluster of phosphorylated Y493 ZAP-70 (red) and expression of DLG1 (blue) In immunostained wildtype or DLG1 KO Jurkat cells. Cells were dropped onto coverslips coated with anti-CD3 (stimulated) and visualized 5 min post stimulation.

D and E. Phosphorylated Y493 ZAP-70 and phosphorylated TCR-ζ intensities were quantified using Image J in immunostained wildtype or DLG1 KO Jurkat cells. Cells were dropped onto coverslips coated with anti-CD3 + anti-CD28 (stimulated) or anti-CD45 (unstimulated). Data representative of 8 fields from 3 independent experiments.

**Supplementary Figure 4** SHP2 knockdown fails to rescue the inhibition of T cell signaling conferred by DLG1 deficiency

A. Immunoblot analysis of lysates from wildtype and DLG1 KO Jurkat cells. Cells were pretreated with control or SHP2 specific siRNA, then activated with soluble anti-

CD3 (monoclonal antibody OKT3) and antibody to mouse immunoglobulin G (IgG), followed by probing for phosphorylated proteins. Membranes were stripped and then reprobed for detection of total cellular proteins corresponding to the phosphorylated proteins.

B and C. Phosphorylated Y493 ZAP-70 and phosphorylated TCR-ζ intensities were quantified using Image J in immunostained wildtype or DLG1 KO Jurkat cells. Cells were pretreated with control or SHP1-specific siRNA, then dropped onto coverslips coated with anti-CD3 + anti-CD28 (stimulated) or anti-CD45 (unstimulated). Data are representative of 8 fields from 3 independent experiments.
